# Vibrissae-evoked activity in the somatosensory thalamus of rat pups: an intracellular study

**DOI:** 10.1101/2022.06.15.496095

**Authors:** Maxim Sheroziya, Roustem Khazipov

## Abstract

Spontaneous and sensory-evoked neuronal activity plays a decisive role in network formation during postnatal development. The thalamus is a major gateway for sensory outputs to the cortex, so that thalamic neuronal activity in newborn animals might be crucial for maturation of the thalamocortical network. The sensory-evoked intracellular thalamic activity and signal propagation in newborn animals remain largely unknown. Here we performed local field potential (LFP), juxtacellular, and patch clamp recordings in the somatosensory thalamus of urethane anesthetized rats at postnatal days 6-7 (P6-7, both sexes) with one whisker stimulation. To reach the thalamus with the electrodes the majority of the overlying cortex and hippocampus were removed. Deflection of only one (the principal) whisker induced spikes in a particular thalamic cell. Sensory stimulation evoked excitatory and inhibitory postsynaptic events in thalamocortical cells. Up to 5-10 sensory-evoked large-amplitude excitatory events followed with 100-200 ms inter-event intervals, while multiple inhibitory events tended to form 20-40 ms inter-event intervals. Large-amplitude excitatory events produced spike bursts with an intraburst frequency of 50-100 Hz and/or short plateau potentials in thalamocortical cells. Inhibitory events could down-modulate evoked spiking or prevented a depolarization block. Juxtacellular recordings confirmed the partial inactivation of spikes during short plateau potentials. Excitatory events evoked low-threshold spikes (LTS) in thalamocortical cells, but, in agreement with previously reported results, hyperpolarizing current pulses generated weak LTS without spike bursts. We conclude that thalamic neuronal activity in rat pups is determined by relatively weak and slow intrinsic membrane currents and relatively strong synapses that might underlay immature forms of thalamocortical synchrony and signal propagation.

## Introduction

Sensory-evoked and spontaneous neuronal activity plays an important role in network formation during development (Blankenship and Feller, 2010; Erzurumlu and Gaspar, 2012; Vitali and Jabaudon, 2014). In line with this basic rule, newborn rodents already have functional vertical somatosensory pathways that are able to transmit sensory-evoked signals to the cortex, which serve network maturation rather than sensation (Khazipov and Milh, 2018; Luhmann and Khazipov, 2018). During the first postnatal week in rodents, the cortex and thalamus generate short-term spontaneous and sensory-evoked gamma oscillations and early alpha/beta oscillations called spindle-bursts, but remain silent most of the time at the LFP level (Minlebaev et al., 2007; Marcano-Reik and Blumberg, 2008; Yang et al., 2009; Minlebaev et al., 2011; Yang et al., 2013). It was proposed that the thalamus and/or periphery might contribute to both early gamma activity and spindle-bursts (Khazipov and Milh, 2018; Murata and Colonnese, 2019). The rodent cortex starts generating global adult-like patterns, i.e. persistent network activity during wakefulness and slow oscillations during sleep and general anesthesia, by approximately the end of the second week of postnatal development (Golshani et al., 2009; Murata and Colonnese, 2018; Dominguez et al., 2021).

The thalamus is the main gateway for sensory signals on their route to the cortex (Hwang et al., 2017). Thalamocortical cells of the relay thalamic nuclei recieve sensory signals via large excitatory synapses (thalamic drivers) formed on proximal dendrites. Feedback corticothalamic axons form small synapses (thalamic modulators) on distal dendrites of thalamocortical cells in the relay nuclei. Feedback corticothalamic axons originating mainly from cortical layer V pyramidal neurons also form drivers in the higher-order nuclei (Sherman and Guillery, 1996, 1998; Groh et al., 2014). Large amplitude and/or fast rising excitatory postsynaptic potentials recorded in thalamic cells in vivo are thought to be generated by thalamic drivers (Brecht and Sakmann, 2002; Deschenes et al., 2003; Ganmor et al., 2010; Sheroziya and Timofeev, 2014, 2015; Urbain et al., 2015; Mease et al., 2016). It has been shown on thalamic slices of newborn animals that NMDA receptors participate strongly in retinogeniculate and somatosensory relay synapses (Ramoa and McCormick, 1994b; Arsenault and Zhang, 2006), but thalamic driver activity has never been described in newborn animals in vivo.

The low-threshold calcium T-current largely defines the intrinsic property of thalamic cells to produce LTS bursts (Jahnsen and Llinas, 1984). Generation of LTS bursts is mechanistically bound to fast transitions from a hyperpolarized to depolarized membrane potential, which is a feature of slow-wave sleep and general anesthesia (Steriade et al., 1993; Crunelli et al., 2006; Llinas and Steriade, 2006; Fernandez and Luthi, 2020). In particular, thalamocortical and inhibitory cells in the reticular thalamic nucleus (TRN) generate rhythmic LTS bursts during thalamic sleep spindles. On the other hand, thalamic drivers produce either spike bursts or unitary spikes (bursting and tonic mode) in a membrane potential dependent manner in thalamic relay cells (Castro-Alamancos, 2002). Behavioral state and behavior itself determine the thalamic firing mode in adult animals (Hirsch et al., 1983; Guido and Weyand, 1995; Ramcharan et al., 2000; Fanselow et al., 2001; Sherman, 2001; Marlinski and Beloozerova, 2014; Urbain et al., 2019). In thalamic slices of newborn animals, neurons generated weak LTS without a spike burst or with a burst of lower spike frequency and generated broad action potentials compared to mature activity (Ramoa and McCormick, 1994a; Perez Velazquez and Carlen, 1996; Pirchio et al., 1997; Tennigkeit et al., 1998; Lee et al., 2010). Weak intrathalamic inhibitory feedback from TRN is already present in newborn rodents (Warren and Jones, 1997; Evrard and Ropert, 2009; Lee et al., 2010). In ferret slices of the lateral geniculate nucleus, spindle oscillations appeared at 3-4 weeks of age (McCormick et al., 1995).

In this study we tested early sensory-evoked activities in the thalamus of urethane anesthetized rat pups. Corticothalamic connections were inactive because we removed the overlying somatosensory cortex. Sensory-evoked rhythmic large-amplitude EPSPs (likely thalamic drivers) produced spike bursts in thalamocortical cells. Membrane potential depolarization did not prevent bursting activity, while IPSPs, likely originating from TRN, could down-modulate bursting firing to the gamma frequency range. Thus, sensory-evoked large-amplitude EPSPs in thalamocortical cells is the main driving force for early thalamic gamma oscillations.

## Methods

### Ethical Approval

Experiments were performed in accordance with EU Directive 2010/63/EU for animal experiments, and the animal-use protocols were in line with Kazan Federal University on the use of laboratory animals (ethical approval by the Institutional Animal Care and Use Committee of Kazan State Medical University N9-2013).

### Surgery

Rat pups (6-7 days old, P6-7) of both sexes were used in this study. Surgery was performed under 2-4% isoflurane anesthesia, and urethane (1-1.3 g/kg, i.p.) was injected by the end of surgery. Animals were placed in a short tube, put on a warm thermal pad (37°C) and carefully covered with cottonwool. Animals were placed in a sitting fetal position in the tube so that the head was bent forward relative to the body. The sitting fetal position allowed the fourth ventricle to be opened (see below) in rat pups which have a short neck. The skull of the animal was cleaned of skin and the head was fixed in the stereotaxic frame with metal plates, dental cement and superglue. A large cranial window was drilled out above the left hemisphere (from bregma to lambda and from 0.2-0.3 mm to 5-6 mm lateral from a midline). Using the cranial window all accessible cortex and the hippocampus (partly) were gently removed with suction (see Fig.7). In 3 experiments with silicon probes (see Fig.2) we drilled out all accessible upper skull bones and removed all upper cortex and hippocampus (lateral: from 0.2-0.3 mm to 4-6 mm from a midline; anteroposterior: from −3 mm to lambda). The obtained recess was covered with paraffin in mineral oil (1/3) during thalamic recordings. To reduce brain pressure and pulsation, we opened the fourth ventricle after drilling. As a reference electrode we used a chlorinated silver wire placed on the cerebellum.

### Extracellular and patch clamp thalamic recordings

All recordings were obtained in the left hemisphere. Thalamic local field potential (LFP) recordings were obtained with linear silicon probes (50 μm spacing between the recording sites, NeuroNexus Technologies, USA). To visualize traces the probes were covered with fluorescent (green light) dye (1,1’-dioctadecyl-3,3,3’,3’-tetramethylindocarbocyanine perchlorate, Sigma). LFP signals from the probes were acquired using a Digital Lynx amplifier (Neuralynx, United States), digitized at 32 kHz, and saved for post hoc analysis. For juxtacellular recordings we used tungsten needle microelectrodes (tip resistance 4-5 MΩ, Microelectrodes, Cambridge, UK). Using a micromanipulator (Sutter Instrument) we slowly immersed electrodes into the thalamus to a depth of ~ 1-1.5 mm (Khazipov et al., 2015) and located vibrissae-evoked responses. Similarly to the adult thalamus, a particular thalamic cell responded with spikes to deflection of only one whisker (the principal whisker). In each experiment we detected a principal whisker manually and then a plastic tube (1 mm tip diameter) connected to Pneumatic PicoPump (WPI) was fixed ~1 mm from the whisker for air puff stimulation. Similarly to the tungsten electrodes, patch pipettes were slowly inserted with a micromanipulator (Sutter Instrument) to the same depth. For current clamp recordings patch pipettes were backfilled with the following solution (in mM): 144 K-gluconate, 4 KCl, 10 disodium phosphocreatine, 10 HEPES, 4 MgATP, and 0.3 GTP, pH 7.3 (tip resistance ~7 MΩ); for voltage clamp (in mM): 144 Cs-gluconate, 4 CsCl, 10 disodium phosphocreatine, 10 HEPES, 4 MgATP, and 0.3 GTP, pH 7.3 (tip resistance ~7 MΩ) with or without 5 mM QX-314 (bromide). Series resistance was 30–60 MΩ. To visualize thalamic cells the pipettes were also filled with 0.5% biocytin (Sigma). Juxtacellular and intracellular patch clamp signals were acquired with a MultiClamp 700B amplifier (Molecular Devices), digitized at 32 kHz and saved for post hoc analysis.

Sensory stimulation was performed with 200-ms-long air puffs every 10 s. In each experiment we positioned the air puff tube to deflect only one whisker on the right side of the snout under visual control. The intensity of air puffs was adjusted by increasing/decreasing air pressure (range 0-10 Psi) with the Pneumatic PicoPump. Air puffs led to ~1 mm whisker deflection and high frequency vibration (~300 Hz, detected with a high speed camera, not shown). To record stable sensory-evoked responses we used an air puff intensity (pressure) 2-3 times higher than the minimal intensity which was able to produce a thalamic response (Fig. 3). Using a piezoelectric sensor we measured the delay (12-13 ms) between actuation of the Pneumatic PicoPump and the onset of the air puff at the other end of the tube (Fig.1). To calculate true response latency we subtracted 12.5 ms from the measured latency.

**Figure 1.**
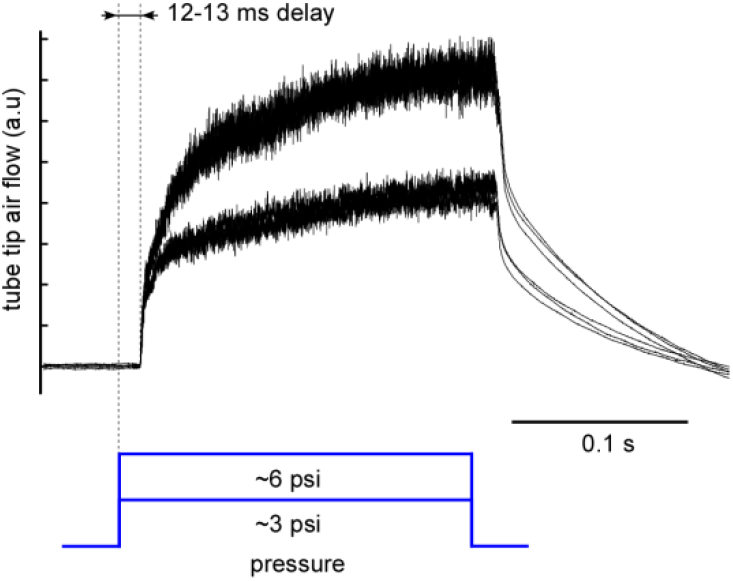
Air puff calibration. A computer controlled Pneumatic PicoPump provided air pressure in a plastic tube (length ~2 m) used for whisker stimulation. Air flow produced by the other end of the tube was detected with a small piezoelectric sensor. Air flow reached the other end of the tube with a 12-13 ms delay after the computer command. The piezoelectric sensor detected air flow as a high frequency noise, while the slow sensor readings represent the sensor deformation and/or offset.

### Histochemistry

After recordings deeply anesthetized animals were perfused with 0.1 M phosphate buffered saline (PBS) followed by 4% PFA in PBS. The brains were then removed and stored overnight in 4% PFA in PBS at 4°C. Then 70-μm-thick frontal sections were obtained with vibratome (Microm HM 650V, Thermo Scientific), extensively rinsed and incubated in 0.3% H_2_O_2_ in PBS for 2 h. To visualize biocytin-labeled cells, free-floating sections were incubated in ABC kit (1:200; Vector Laboratories), 0.1% Triton X-100 (Panreac) and 1% bovine serum albumin in PBS for 3 h. After rinsing, sections were incubated in PBS containing 0.05% DAB (Sigma) and 0.00125% H_2_O_2_ for 5–10 min.

### Data Analysis and Statistics

Signals were analyzed using built-in functions and custom-developed MATLAB scripts (MathWorks, United States). To detect multiunit activity (MUA) in LFP recordings (silicon probes), we high-passed (100-15999 Hz) the raw wide-band signal, differentiated the signal, calculated standard deviation (SD) and then located negative peaks below a threshold of 4-5 SD. The detected MUA were averaged and visually inspected. For spike detection in juxtacellular recordings we used a threshold of 2-4 mV for the raw signals with visual inspection. For each extracellular recording we plotted a peristimulus time histogram and detected latency (the first bin with a value above zero) for sensory-evoked spikes. For spectrograms (Figs. 5 and 7) we re-sampled the raw signals from 32 to 1 kHz, set the baseline to zero, calculated a spectrogram for each sensory-evoked response (Hamming window 328 ms), and averaged the spectrograms. In order to produce similar and comparable spectral analysis for MUA activity obtained with silicon probes (Fig.4) we used a high-pass filter (>1.5 kHz) and smoothed LFP signals; then we calculated the spectrograms as described above. In order to compare spectrums of intracellular sensory-evoked currents (Fig.7) we re-sampled the raw signals from 32 to 1 kHz, set the baseline to zero, normalized each response on its mean absolute deviation and then calculated the spectrum. To detect significant differences we employed the nonparametric Wilcoxon signed rank test. To detect IPSCs (Fig.7) we differentiated the signals, calculated SD and located positive peaks above 2 SD. For EPSCs we differentiated the raw signals and located negative peaks below 32 pA/ms. In the figures the data are expressed as mean value ± SEM. In the text the data are expressed as mean value ± SD.

## Results

### LFP and juxtacellular thalamic responses to one-whisker stimulation in rat pups

We recorded vibrissae-evoked responses in the thalamic ventral posteromedial nucleus (VPM) using linear silicon probes with 50 μm spacing between the recording sites. The silicon probes were covered with fluorescent dye and insertion traces were located after recordings. We removed all accessible upper cortex and partially hippocampus before all recordings (see methods), so that the barrel cortex could not provide feedback to the thalamus during the sensory response. In the experiments with silicon probes we removed almost all the upper part of the left hemisphere (Fig.2). Averaged LFP responses composed of a short-term ripple or gamma oscillation followed by a long (up to 1 sec) slow negative shift were detectable in 3-4 LFP channels of the probe. The amplitude of the slow negative shift varied from 30 μV to 180 μV in our recordings (n = 3 rats). In the shown example (Fig.2), we successively recorded LFP responses from 2 neighboring whiskers in a given location of the probe. Air puff stimulation of the first vibrissae (C3) produced responses predominantly in channels 3-5, while the second vibrissae (D2) evoked responses mainly in channels 6-8 (the channels at the tip of the silicon probe). Stimulation of other whiskers did not evoke responses in the shown location (Fig.2) of the silicon probe. We detected MUA for each LFP channel and plotted averaged peristimulus time histograms which revealed similar depth-profile to sensory responses. In 2 other recordings with silicon probes (not shown) MUA responses covered 4 channels of the probe with 50 μm spacing between the recording sites. Thus the detected vertical size of thalamic barreloids was 150-200 μm. On average, 1-2 silicon probe channels with stronger responses displayed shorter response latency and 1-2 other channels showed longer latency: with the minimal latency 23 ± 1 ms and maximal 28 ± 1 ms. The duration of the responses varied from 1 to 5 s. In the present study we used 200-ms-long air puffs for one-whisker stimulation. Strictly speaking, air flow could affect neighboring whiskers located on in-line with the air puff tube, but since we were able to detect the bounds of thalamic barreloids, the air puff stimulation was local enough for our purposes.

**Figure 2.**
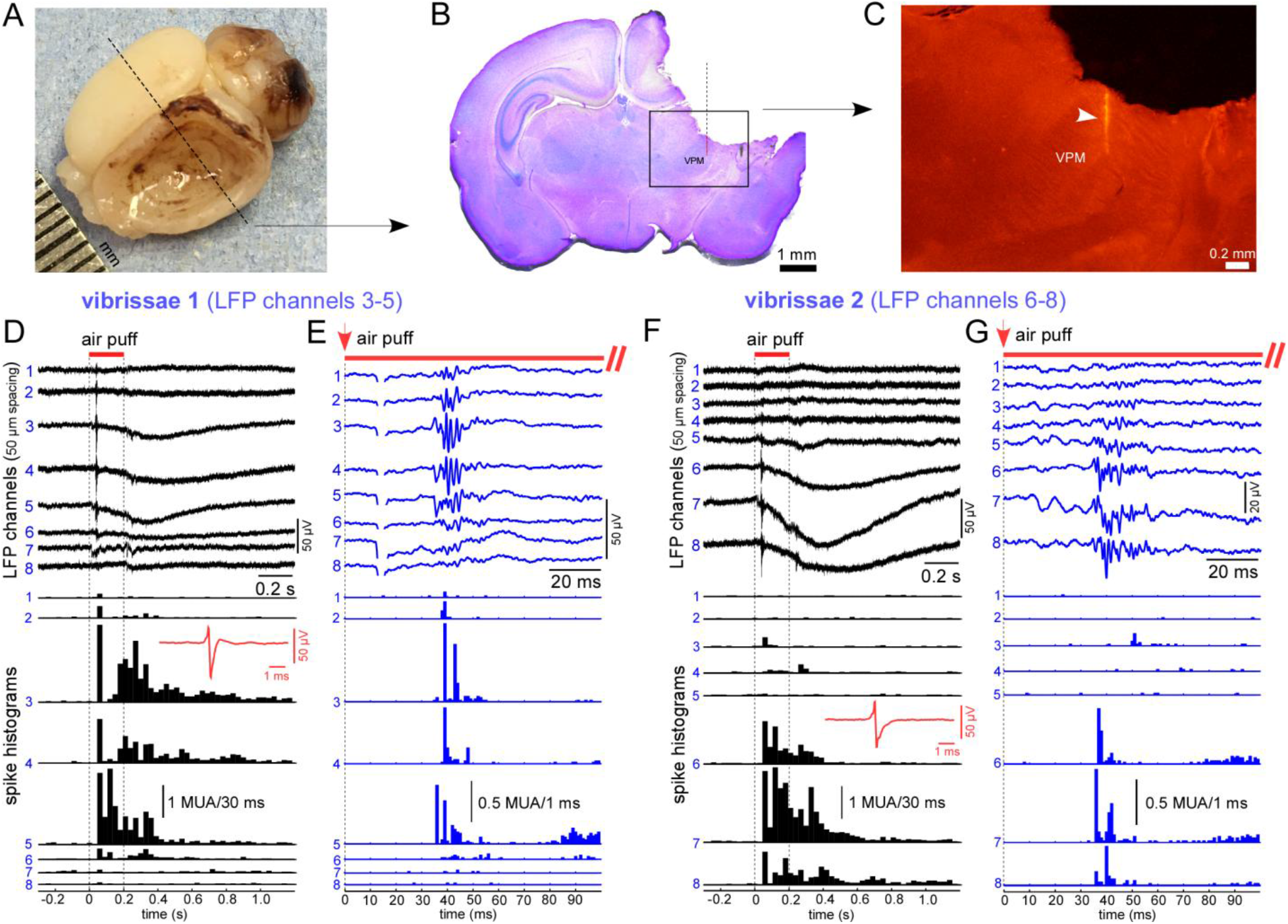
Thalamic sensory-evoked responses recorded with a linear silicon probe (low impedance LFP electrodes). (A) A photo of a formalin fixed rat brain. In order to reach the thalamus with electrodes the majority of upper cortical areas and hippocampus in the left hemisphere were gently removed. The dashed line schematically indicates the location of the frontal section shown in B. (B) A light photograph shows the frontal section stained with cresyl violet. (C) A fluorescent trace of the silicon probe in the thalamus. The trace of the silicon probe covered with a fluorescent dye was detected in green light before staining with cresyl violet. (D) Averaged (60 responses, vibrissae 1) one whisker-evoked LFP responses with peristimulus time histograms (below) recorded in the brain shown in A, B and C with a linear silicon probe (8 of 16 channels shown, 50 μm spacing between recording sites). Channel numbers are shown in blue. The red line represents the air puff (duration 0.2 s). Peristimulus time histograms represent MUA spiking of the corresponding LFP channels (see channel numbers). An example of an averaged MUA spike is shown in red. (E) The onset of the same sensory-evoked responses shown in D. The red arrow represents air puff onset. (F) Averaged (60 responses) LFP responses with peristimulus time histograms (below) evoked with an air puff stimulation of a neighboring vibrissae. An example of averaged MUA spike is shown in red. Note that sensory-evoked responses were longer than stimulus duration. (G) Onset of the sensory-evoked responses shown in F. Red arrow represents air puff onset.

The duration and amplitude of sensory-evoked thalamic responses depended on air puff intensity. In Figure 3 we show LFP and juxtacellular thalamic responses evoked with weaker and stronger air puff stimulation. Stronger stimuli evoked pronounced bursting thalamic activity lasting about 5 s in the example shown with juxtacellular recording (Fig.3C). In all other experiments we used an air puff intensity 2-3 times higher than the minimal intensity that produced a thalamic response. Air puff stimulation evoked deflection and fast whisker vibration/fluttering in all directions because of turbulence. For this reason air puff stimulation probably provides the strongest whisker stimulation.

**Figure 3.**
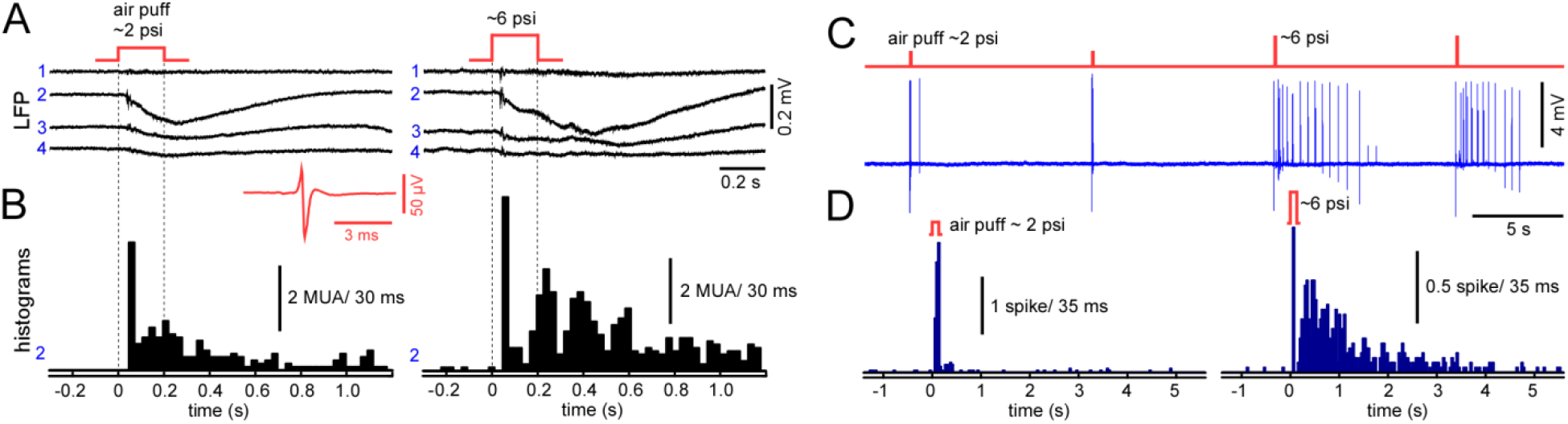
Thalamic one whisker-evoked response depended on air puff intensity. (A) Averaged (10 responses) LFP responses evoked with weak (left) and stronger (right) air puff stimulation (4 of 16 silicon probe channels shown, 50 μm spacing between recording sites). Channel numbers are shown in blue. (B) Peristimulus time histograms plotted for channel 2 shown in A. An example of averaged MUA spike is shown in red. (C) Juxtacellular sensory-evoked thalamic responses recorded with a high impedance (4-6 MΩ) extracellular electrode during weak (left) and stronger (right) air puff stimulation. (D) Peristimulus time histograms (30 juxtacellular responses averaged) for weak (left) and stronger (right) air puff stimulation.

To reveal rhythmic MUA during sensory-evoked thalamic responses we filtered (high-pass > 1.5 kHz) and smoothed raw signals. With this manipulation we transformed MUA spikes into a wave and calculated spectrograms (Fig.4). MUA recorded with a silicon probe produced ‘hot spots’ within the gamma and beta frequency ranges in spectrograms (Fig.4C).

**Figure 4.**
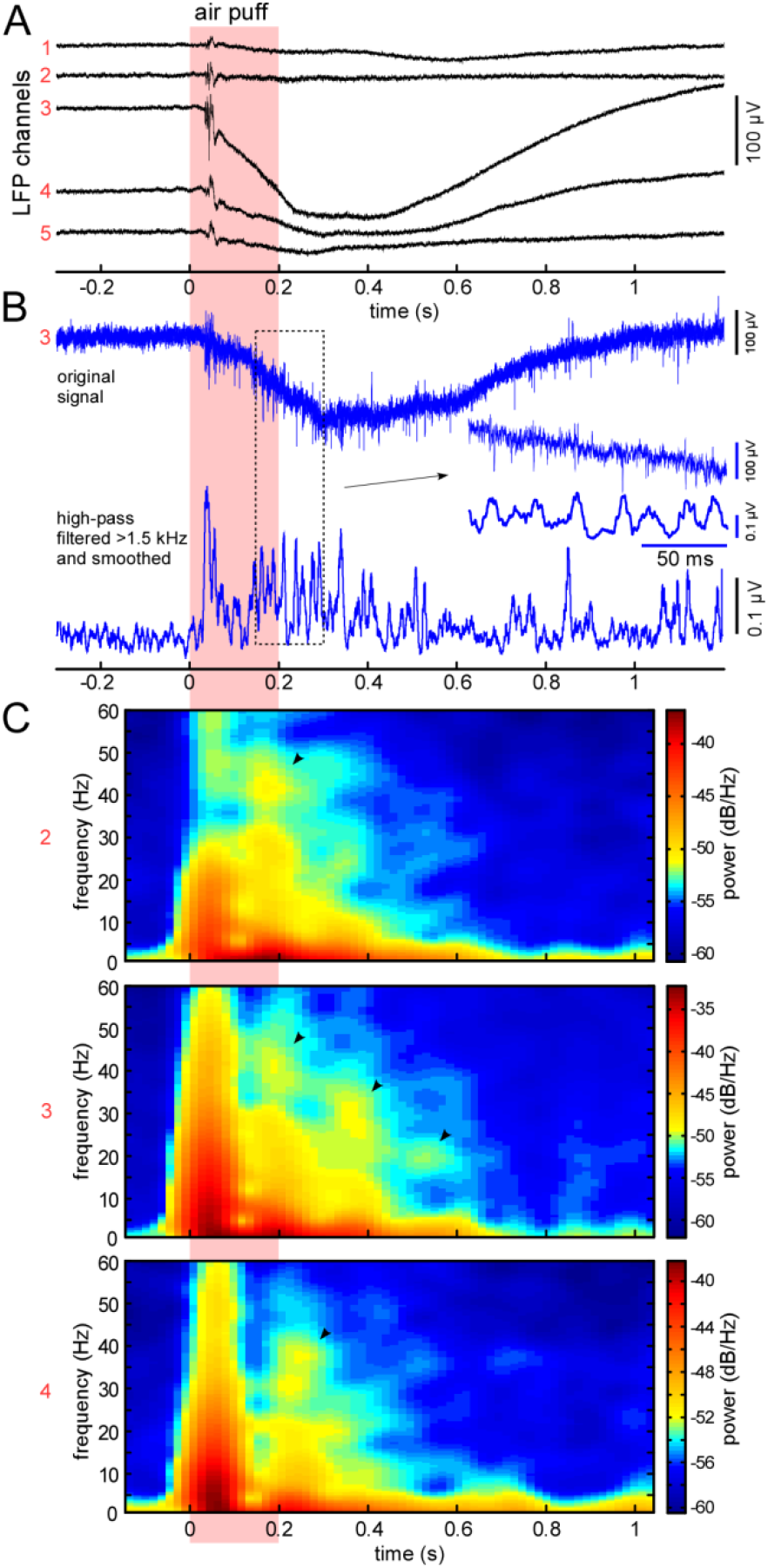
Sensory-evoked thalamic MUA displayed gamma/beta rhythms. (A) Averaged (60 responses) sensory-evoked LFP response recorded with a linear silicon probe (5 of 16 channels shown, 50 μm spacing between recording sites). Channel numbers are shown in red on the left. (B) An example of one thalamic response obtained with LFP channel 3 (raw signal) and high-pass filtered (>1.5 kHz) and smoothed signal below. The filtered and smoothed signal displayed oscillations within the gamma/beta frequency range. (C) Averaged spectrograms calculated for high-pass filtered and smoothed responses for channels 2, 3 and 4 shown in A. “Hot spots” within gamma/beta frequency range are marked with black arrows.

In order to investigate sensory-evoked spiking activity in the thalamus of rat pups we recorded juxtacellular responses (n = 31 recordings in 16 pups). The electrodes used (needle tungsten electrodes, tip size ~10 μm) allowed us to record extracellular ~ 5 mV amplitude spikes from 1-2 thalamic cells (large-amplitude spikes were likely generated by 1 cell). In all experiments thalamic cells responded with spikes to deflection of only one (the principal) whisker. Sensory stimulation evoked a series of spike bursts or groups of spikes and unitary spikes which tended to form 100-200 ms intervals (Fig.5). We detected spikes and plotted time peristimulus histograms. The latency of spike responses was 29.9 ± 4.2 ms. Averaged stereotypic responses formed an oscillation-like pattern within the alpha frequency range (Fig.5B). We also calculated the distributions of inter-spike intervals (ISIs) for sensory-evoked thalamic activity. The distributions displayed 2 peaks corresponding to intra-burst and inter-burst intervals (Fig.5C) in 14 (~45%) juxtacellular recordings. Peak locations of intra-burst and inter-burst ISI distributions were 13.5 ± 2.3 ms and 118 ± 34 ms (range 83-205 ms) respectively. Compared to the adult thalamus intra-burst spike frequency in pups was smaller and bursts did not show an ISI increase with ISI ordinal in a given burst (examples in Fig.5C), a known feature of LTS bursts in thalamic cells (Lenz et al., 1994; Jeanmonod et al., 1996; Wei et al., 2011). Instead, spike bursts could be produced by strong excitatory postsynaptic events (see below).

**Figure 5.**
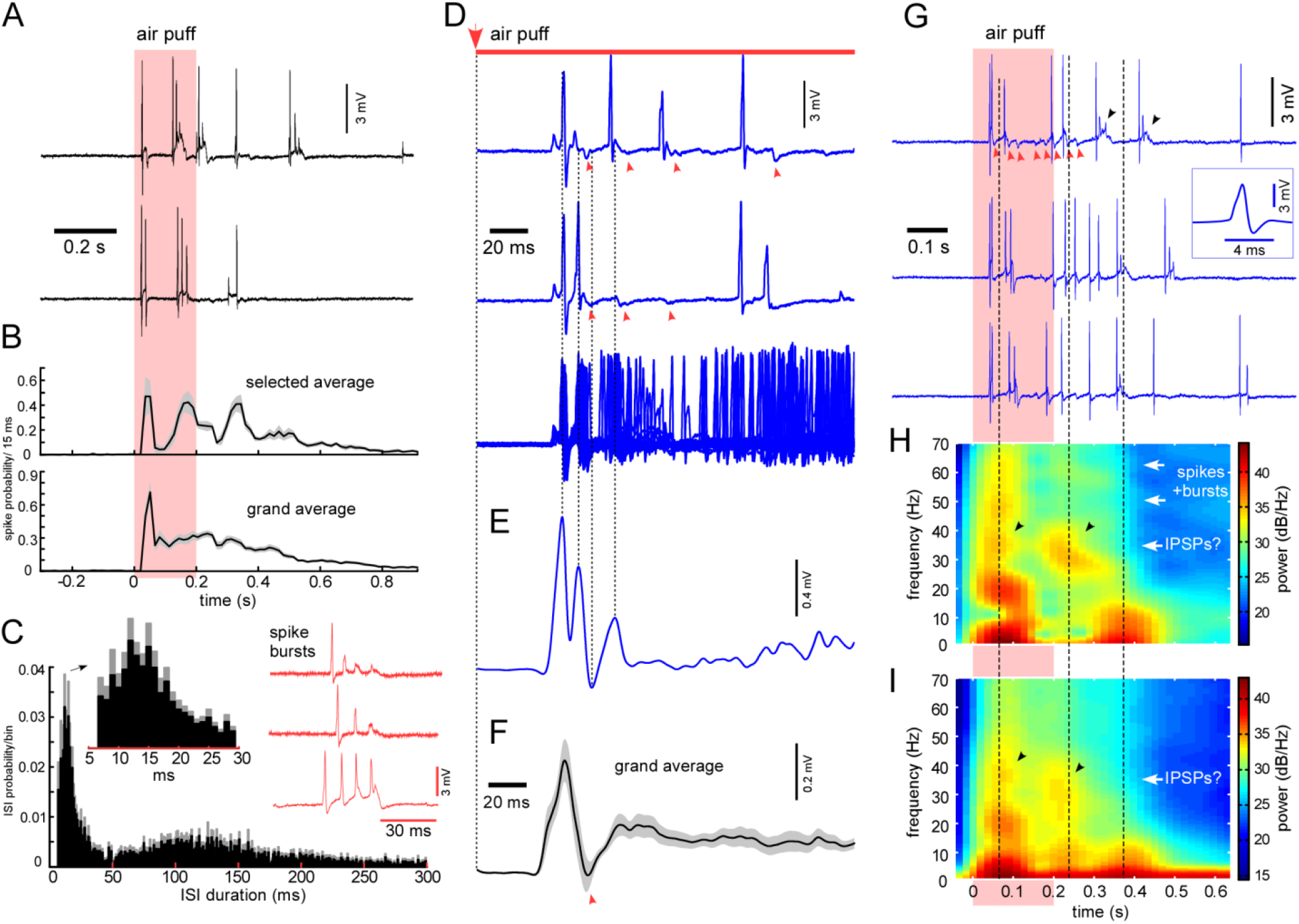
Thalamic juxtacellular sensory-evoked responses recorded with a high impedance extracellular electrode. (A) Two examples of sensory-evoked responses. One-whisker stimulation evoked a series of spike bursts in thalamic cells. Air puff stimulation is shown in pink. (B) Averaged peristimulus time histograms represent thalamic spiking response. A majority of thalamic cells displayed tendency to fire in bursts every 100-200 ms after sensory stimulation. The upper histogram shows the averaged response for 9 recordings which displayed similar inter-burst intervals. The bottom histogram shows the grand average sensory-evoked response (n = 31 recordings, 60 responses each recording). (C) Bimodal distribution of inter-spike intervals (ISIs) during sensory-evoked response. In 14 (~45%) juxtacellular recordings ISI distribution displayed 2 peaks corresponding to intra-burst and inter-burst intervals. An example of a spike burst is shown in red. (D) Two examples of the onset of the sensory-evoked spiking responses and a superposition of 60 responses below. Note negative peaks are indicated with red arrows. (E) Spiking responses shown in D form oscillation after averaging. The responses were smoothed with a low-pass filter (100 Hz) and averaged. (F) Grand average thalamic spiking response (n = 31, 60 responses each recording). (G) Gamma-modulated thalamic spiking activity during sensory-evoked response. Three examples of sensory responses modulated in the gamma frequency range. In the first example the negative peaks are shown with red arrows. Note that likely the same thalamic cell could generate spike bursts (shown with black arrows). An example of averaged spike is shown in a frame. (H) Averaged spectrogram calculated for 60 responses for the juxtacellular recording shown in G. Hot spots within gamma frequency range marked with black arrows. (I) Averaged spectrogram for 7 recordings (60 responses each) which displayed similar negative peaks (see D) and gamma hot spots (marked with black arrows).

Stereotypic neuronal spiking at the beginning of the responses led to oscillation-like patterns after averaging (Fig.5E). In 7 recordings we distinguished low-amplitude negative peaks which are marked with red arrows in Fig.5. Since the negative peaks were in opposition to spikes, they may represent intracellular hyperpolarizing and inhibitory events (see below). Rhythmic low-amplitude negative peaks produced oscillatory patterns within the gamma frequency range during sensory responses (Fig.5G). To reveal evoked gamma activity we re-sampled raw signals to 1 kHz, calculated a spectrogram for each response and then plotted an average spectrogram (Fig.5H). Low-amplitude negative peaks produced hot spots within the gamma frequency range in spectrograms (marked with black arrows in Figs.5H and I). Based on the obtained extracellular data, we propose that sensory stimulation evoked a series of relatively strong excitatory postsynaptic events able to produce spike bursts which, in turn, are gamma-modulated with inhibitory events.

In one juxtacellular recording we observed spontaneous transitions between tonic and bursting firing (the only recording with spontaneous bursting). Bursts displayed ISI increase with ISI ordinal in a given burst (Fig.6), a known feature of LTS bursts (Lenz et al., 1994; Jeanmonod et al., 1996).

**Figure 6.**
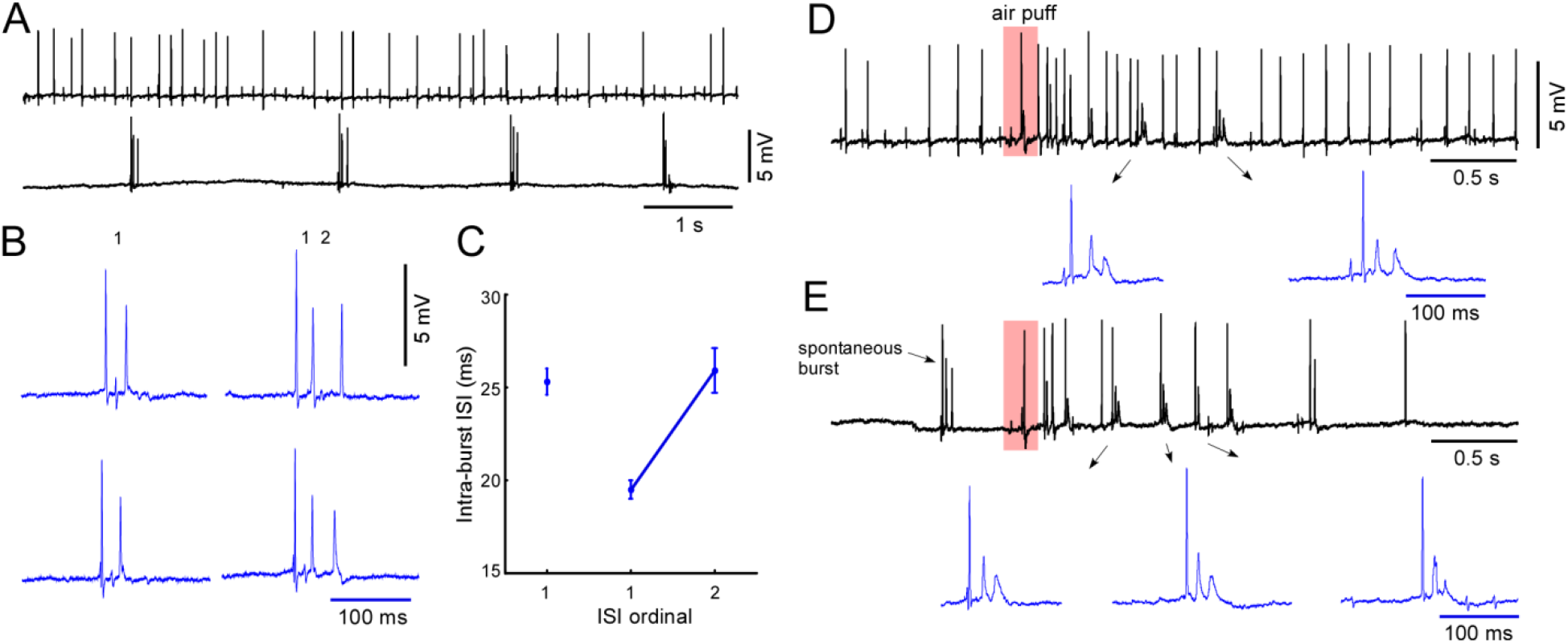
Spontaneous tonic and bursting firing of thalamic cells. (A) Cellular activity spontaneously switched between tonic and bursting firing (corresponding examples are shown; juxtacellular recording). (B) Examples of spontaneous spike bursts. (C) Averaged intra-burst ISIs. Spontaneous bursting activity displayed a known feature of LTS bursts. (D) Sensory-evoked response during spontaneous tonic firing. Examples of evoked spike bursts are shown in blue. (E) Sensory-evoked response during spontaneous bursting firing. Examples of evoked spike bursts are shown in blue. Decrease of spike amplitudes in sensory-evoked spike bursts indicated strong depolarization and partial inactivation (depolarization block) of fast sodium channels.

### Patch clamp recordings of sensory-evoked responses in the thalamus of rat pups

To test our hypothesis that thalamic sensory responses composed of excitatory postsynaptic events are modulated with inhibition we performed patch clamp recordings. As above, we removed the upper cortex and partially hippocampus, therefore corticothalamic inputs did not affect thalamic responses. Immediately after break-in we applied current steps in current clamp mode. Thalamic cells generated LTS without spike bursts after a hyperpolarizing current step (Fig.7E). We performed voltage clamp recordings with Cs-gluconate-based solution with and without QX-314 (n = 9 cells in 7 pups). We could clamp a membrane potential in a range from ~ 0 mV to ~ −70 mV (Fig.7A); though in the experiments without QX-314 spikes sporadically occurred at hyperpolarized potentials (not shown). Cell input resistance was 0.46 ± 0.07 GΩ. Thalamic sensory-evoked response composed of a series of excitatory postsynaptic currents (EPSCs) distinguished at hyperpolarized potentials and inhibitory postsynaptic currents (IPSCs) manifested at depolarized membrane potentials (Fig.7A). The mean number of EPSCs per response was 5.13 ± 1.8 (range 3-9). In the example of biocytin stained thalamocortical cell shown (Fig.7D), the reconstructed axon of the cell gives collateral in TRN. The latter indirectly confirms that TRN provided inhibitory feedback to VPM during sensory-evoked responses in rat pups. A spectrogram calculated for sensory-evoked responses at depolarized membrane potentials revealed hot spots within the gamma frequency range (Fig.7B), comparable to the gamma activity observed in the juxtacellular recordings. Accordingly, the gamma power of sensory-evoked currents significantly decreased at a membrane potential of −70 mV compared to 0 mV (n = 9 cells; Fig.7C). We also were able to detect EPSCs in a ‘tail’ of sensory-evoked responses and to check their dependency on the holding membrane potential (Fig.7F and G). In agreement with previously reported data (Ramoa and McCormick, 1994b; Arsenault and Zhang, 2006), the nonlinear voltage-current dependency of peak amplitudes and the different shape of EPSCs indicated that NMDA receptor-mediated current contributed strongly to sensory-evoked EPSCs in VPM of rat pups. We detected postsynaptic events, calculated inter-IPSCs and inter-EPSCs intervals, and plotted their distributions. Inter-IPSC distribution displayed a plateau around 20-40 ms intervals (Fig.7I) in contrast to many electrophysiological distributions which have an exponential right tail (Buzsaki and Mizuseki, 2014). The peak location for inter-EPSC interval distribution was 183 ± 85 ms (range 90 – 300 ms). Both distributions show that corresponding sensory-evoked postsynaptic events tended to generate rhythms and were generated either by one (or a small number of) presynapse with its intrinsic frequency, or by many synchronized presynaptic cells.

**Figure 7.**
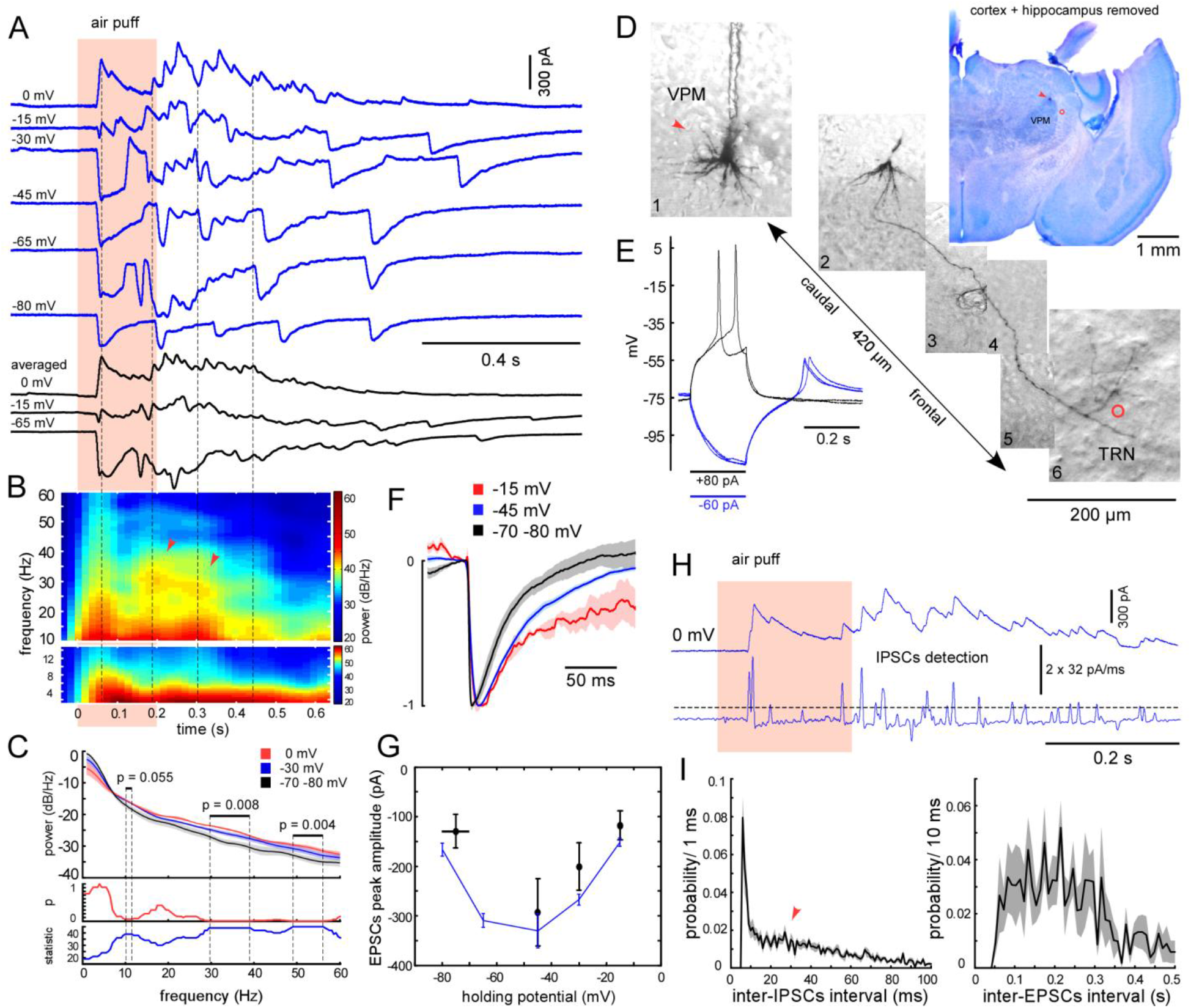
Voltage-clamp recordings from thalamic cells revealed sensory-evoked excitatory and inhibitory postsynaptic currents. (A) Sensory-evoked intracellular currents recorded at different membrane potentials in the biocytin stained thalamocortical cell shown in D. Averaged responses are shown below in black (6-10 responses averaged). Pink area represents air puff stimulation. The recording was performed with Cs-gluconate-based pipette solution with QX-314. (B) Averaged spectrogram calculated for 10 responses recorded at 0 and −15 mV holding membrane potential, one of which is shown in A. Hot spots within gamma frequency range marked with red arrows. Note approximate coincidence of gamma hot spots with large EPSCs (vertical dashed lines). (C) Comparison of the response spectrums at different holding membrane potentials. Each response was normalized on its mean absolute deviation before spectrum calculation (see methods). The upper plots represent averaged spectrums for 3 holding membrane potentials (0, −30 and −70-80 mV, n = 9 cells). The bottom plots display p-value and value of Wilcoxon signed rank test statistic (0 mV vs. −70-80 mV). The gamma power of the sensory-evoked responses was significantly higher at 0 mV compared to hyperpolarized membrane potential. (D) Biocytin stained thalamocortical cell with a reconstructed axon. Cresyl violet stained frontal section is shown in the upper right corner (the biocytin stained cell marked with a red arrow). Combined 6 grayscale images represent the stained cell with its axon which gives collateral in TRN (marked with a red circle). (E) Cellular responses to depolarizing and hyperpolarizing current pulses obtained in current-clamp mode immediately after cell opening. Note LTS without spikes (blue). (F) The shape of sensory-evoked EPSCs depended on the holding membrane potential (grand average, n = 7). (G) Peak amplitude of sensory-evoked EPSCs depended on the holding membrane potential (black, n = 7 cells). EPSC peak amplitudes for the example recording in A are shown in blue. (H) Detection of sensory-evoked IPSCs. Examples of intracellular signal at 0 mV holding membrane potential and differentiated signal with a threshold (dashed line). We detected inter-IPSCs intervals > 5 ms. (I) Averaged distribution of inter-IPSCs (left, n = 9 cells) and inter-EPSCs (right, n = 7 cells) intervals during sensory-evoked responses. Liquid junction potential correction (15 mV subtraction) was applied to all scales.

To confirm our findings we performed current clamp recordings with K-gluconate-based pipette solution (n = 7 cells in 7 pups, Fig.8). Thalamic cells generated LTS either without spikes or with one spike after a hyperpolarizing current step and the input resistance was 0.44 ± 0.09 GΩ. Whisker deflection evoked a series of spike bursts/plateau potentials modulated with inhibitory potentials. To compare sensory-evoked responses at different membrane potentials we applied constant current injections. Large-amplitude excitatory postsynaptic potentials (EPSPs) were able to produce LTS in a membrane potential-dependent manner (Fig.8B and D). Similarly to extracellular recordings, the ISI distribution of intracellular sensory-evoked activities displayed 2 peaks corresponding to intra- and inter-bursts intervals (Fig.8F). Peak locations of intra-burst and inter-burst ISI distributions were 12.7 ± 1.8 ms and 108 ± 12 ms respectively. The intra-burst peak was more pronounced for intracellular recordings (peak amplitude ~0.5 probability/1ms) compared to the intra-burst peak for juxtacellular recordings (peak amplitude ~0.3 probability/1ms, Fig.5C) because spike amplitudes in a given extracellular burst decreased with number of spikes in a burst and were missed with constant threshold detection (see methods).

**Figure 8.**
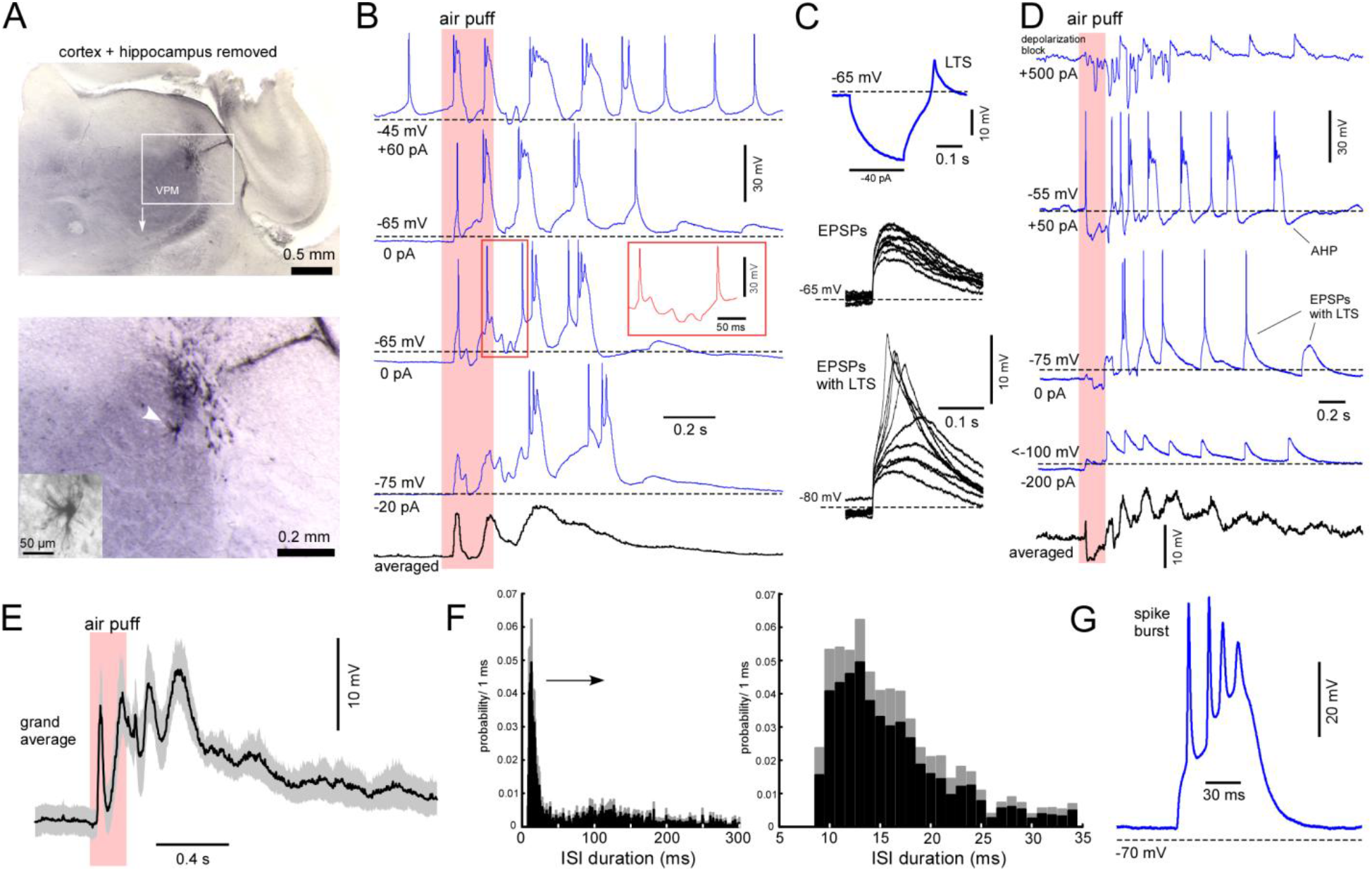
Sensory-evoked large-amplitude EPSPs produce spike bursts modulated with IPSPs. (A) Frontal section with a thalamocortical cell stained with biocytin during patch clamp recording. Positive pressure in the patch pipette on its way to VPM led to extracellular staining with spilled biocytin. The cell was approached with zero pipette pressure which explains the gap between the cell (marked with an arrow) and the dark area. (B) Examples of sensory-evoked responses of the cell shown in A. To change membrane potential we applied constant current injections (indicated next to the examples). Note modulation by IPSPs shown in the red frame. An averaged response is shown below in black (60 responses). (C) Thalamic cells generated rebound LTS after negative current pulse without a spike burst (blue, same cell as in A and B). Large-amplitude EPSPs could generate LTS in a membrane potential-dependent manner (superposition of selected EPSPs from a “tail” of sensory-evoked responses, black, same cell as in A and B). (D) Another example of current-clamp recording from a thalamocortical cell. Depending on the membrane potential sensory-evoked EPSPs produced LTS with single spikes (−75 mV) and spike bursts/plateau potentials (−55 mV), probably enhanced with NMDA currents and/or L-type calcium current (see Discussion). Note the afterhyperpolarization potential (AHP). Mixed excitatory and inhibitory postsynaptic activity at the beginning of the response did not generate rhythmic bursts. (E) Averaged sensory-evoked response recorded in current-clamp mode (grand average, n = 7 cells). (F) ISI distribution of intracellular activities displayed 2 peaks corresponding to intra- and inter-bursts intervals (n = 7 cells). (G) An example of sensory-evoked spike burst. Liquid junction potential correction (15 mV subtraction) was applied to all scales.

To confirm our findings we also performed thalamic recordings (simultaneously with EEG from the barrel cortex) with an intact cortex (Fig.9). We observed similar sensory-evoked activities with juxtacellular electrodes: sensory stimulation produced a series of spike bursts in the thalamus (not shown). Intracellular thalamic activities were obtained with K-acetate-filled sharp electrodes (Fig.9A, B and C). Compared to patch pipettes the sharp electrodes produced shunting effects in immature thalamic cells. The cell input resistance was about 0.1 GΩ (vs. ~0.45 GΩ with patch recordings) and large-amplitude EPSPs produced unitary broad spikes. Cortical sensory-evoked gamma oscillations were detectable at the EEG level and correlated with IPSPs in a VPM cell (Fig. 9B). On the way of the electrodes to VPM we recorded intracellular activities which correlated to cortically generated spindle-bursts (Fig.9C). In adult anesthetized mice, cells in the sensory and higher-order thalamic nuclei display two clearly distinct intracellular patterns: thalamic relay cells follow cortical active/UP states mainly with IPSPs, while neurons in higher-order thalamic nuclei display active-like states. The cells of posteromedial thalamic nucleus (PO), located above VPM, generate active-like states during cortical active/UP states (Sheroziya and Timofeev, 2014; Mease et al., 2016). Thus, putative PO cell displayed active-like state during sensory-evoked cortical spindle-burst (Fig.9C). We also performed LFP recordings in the barrel cortex and calculated spectrograms for sensory-evoked activities (Fig.9D and E). We observed hot spots in gamma frequency range similar to those we observed in thalamic recordings. We propose that the thalamus entrained the cortical cells in graded gamma oscillations (see below).

**Figure 9.**
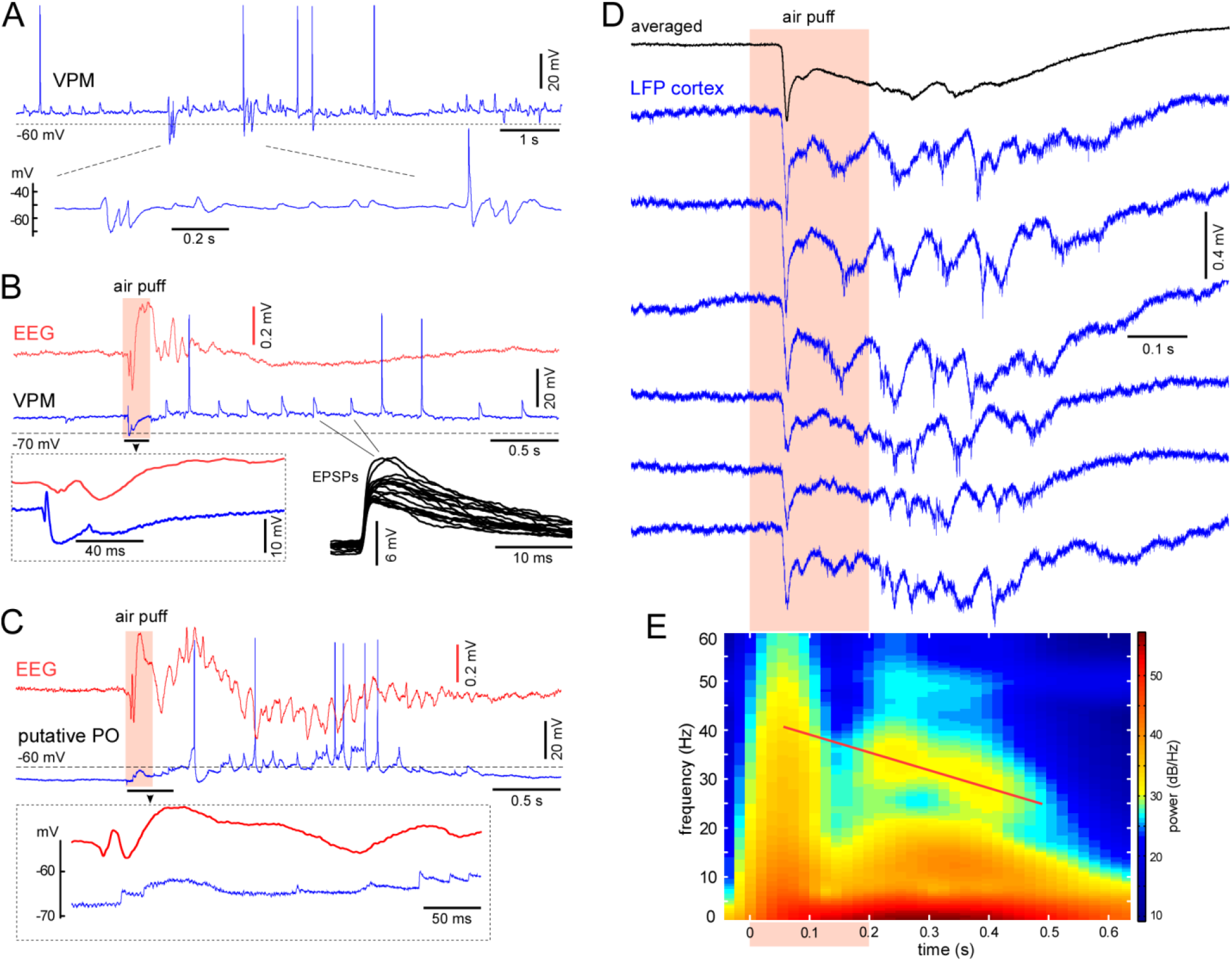
Intracellular thalamic activities with the intact cortex. (A) Spontaneous intracellular activity of VPM cell recorded with a sharp electrode. Spontaneous activity is composed of a constant flow of postsynaptic events. IPSPs and EPSPs tend to form short-term oscillation-like patterns within gamma/beta frequency range. No liquid junction potential correction was used for sharp electrode recordings. (B) Sensory-evoked response of the VPM cell and cortical EEG. Whisker deflection evoked long-lasting response in the VPM cell. Note short-term gamma oscillation with following spindle-burst at EEG. Gamma oscillation correlated with IPSPs in the VPM cell. (C) Sensory-evoked response of a putative PO cell and cortical EEG. Note active-like state in the thalamic cell during cortically generated spindle-burst. (D) Cortical sensory-evoked responses obtained with a needle LFP electrode in the barrel cortex (6 responses are shown with blue). The responses show spindle-bursts and gamma oscillations. (E) Spectrogram calculated for 60 cortical sensory-evoked responses shown in D. Graded gamma activity is schematically marked with a red line.

## Discussion

In the present study we recorded sensory-evoked activities from the thalamic VPM nucleus in rat pups. Deflection of only one whisker induced spikes in a given thalamic cell. At the intracellular level, one-whisker stimulation with an air puff evoked a series of large-amplitude excitatory events modulated with inhibitory events within gamma/beta frequency range. Large-amplitude EPSPs produced weak LTS in a membrane potential dependent manner, spike bursts and plateau potentials. The evoked spike bursts and plateau potentials were generated by a mixture of postsynaptic and intrinsic membrane currents.

### Sensory-evoked postsynaptic events in VPM cells

Thalamic drivers are a known source of large-amplitude excitatory postsynaptic potentials in VPM cells (Brecht and Sakmann, 2002; Castro-Alamancos, 2002). In adult rats sensory-evoked subthreshold responses (EPSPs and IPSPs) in VPM cells ‘could always be elicited from several adjacent whiskers and lasted for hundreds of milliseconds’ (Brecht and Sakmann, 2002), but the response tails did not take much attention so far. Spontaneous postsynaptic activity of many thalamic VPM and LGN cells in adult anesthetized rodents was represented by a persistent flow of large-amplitude EPSPs (Deschenes et al., 2003; Sheroziya and Timofeev, 2014, 2015). Rhythmic EPSPs of prethalamic origin have been reported in the ventroposterior nuclei of the thalamus (Pinault and Deschenes, 1992; Nunez et al., 1994) and it has been reported that whisker-related neurons in the sensory trigeminal complex of adult rats can generate rhythmic activity, in particular, neurons of the nuclei principalis generated a ~10 Hz rhythm (Panetsos and Sanchez-Jimenez, 2010). Importantly, neurons of the trigeminal complex generated LTS and displayed intrinsic oscillatory properties in slices of adult animals and pups (Sandler et al., 1998; Lo et al., 1999; Lo and Erzurumlu, 2001). It is unclear whether electrically coupled thalamocortical cells (reported coupling coefficient at P6-7 was ~0.02) in rodent pups (Lee et al., 2010) can contribute to EPSPs in VPM cells. Thus, according to our current knowledge, the sensory-evoked EPSPs shown here were generated by synaptic inputs from the trigeminal complex. Sensory-evoked EPSPs produced long-lasting spiking responses in our experiments. In adult animals only the subthreshold response lasted hundreds of milliseconds, while the spiking response was restricted to a short time window (Brecht and Sakmann, 2002).

Whisker-specific segregation is already established in VPM cells by P4-5 in rodents (Belford and Killackey, 1979; Yamakado, 1985; Munoz et al., 1999). Developmental remodeling of the sensory inputs involves synapse formation, then local pruning of subsets of axonal branches and synapse elimination (Chen and Regehr, 2000; Arsenault and Zhang, 2006; Takeuchi et al., 2014). In thalamic slices of adult mice a majority of VPM neurons showed an all-or-none EPSC following medial lemniscus stimulation, while in pups thalamic neurons displayed an incremental increase in EPCS amplitude with stronger stimulus intensities. Onsets of EPSCs from different axons were indistinguishable and the number of immature inputs was usually estimated with the number of discrete steps in EPSC amplitude. The number of lemniscal synapses increases during the first postnatal week and decreases after P7-9 (Takeuchi et al., 2014). The mean estimated number of lemniscal axons innervated each VPM neuron was 7-8 at P7-9 and 1-3 in adult rodents (Deschenes et al., 2003; Arsenault and Zhang, 2006; Takeuchi et al., 2014). The maximal amplitude of AMPA-EPSC evoked with medial lemniscus stimulation at −70 mV was ~ 400 pA at P7-9 and ~ 1000 pA after P11-13 (Arsenault and Zhang, 2006). On the other hand, input resistance of thalamic relay cells decreases by 2-3-fold between P6-7 and after P12-14 (Ramoa and McCormick, 1994a; Lee et al., 2010), so that amplitudes of produced postsynaptic potentials (proportional to a product of input resistance and postsynaptic current) would not show dramatic increase with development. In our data, the first sensory-evoked large-amplitude excitatory event was always shunted with inhibition and succeeding large-amplitude excitatory events displayed a shape of unit postsynaptic event. In adult rats firing of trigeminothalamic cells depends on the amplitude, velocity, and direction of vibrissal deflection (Deschenes et al., 2003). Here we used 200-ms-long stimulus and showed that increase in stimulus intensity prolonged the response duration in thalamic VPM cells. Firstly, sensory-evoked poststimulus thalamic activities might directly support early cortical activities (Khazipov et al., 2004). Second, repetition of sensory-evoked signals might serve to developmental maturation of thalamocortical network in newborn animals (Minlebaev et al., 2011).

The electrophysiological impact of recently discovered interneurons in different nuclei of the rodent thalamus (Jager et al., 2021) still remains unclear, thus, the sensory-evoked IPSPs in our experiments likely originated from TRN (Arcelli et al., 1997; Fogerson and Huguenard, 2016). Compared to excitatory response duration of the inhibitory response in VPM cell was shorter presumably because late non-correlated activity of different VPM cells could not activate TRN cells. Due to shorter duration of the inhibitory response we were able to compare effects of mixed postsynaptic events and pure excitatory events (see below).

### Interaction of postsynaptic events and intrinsic membrane currents

In thalamic slices of newborn rats, strong stimulation of the optic tract evoked EPSPs that gave rise to a plateau potential accompanied with a depolarization block of fast sodium channels. Noninvasive juxtacellular recordings confirm partial inactivation of action potentials. Synaptically evoked plateau potential relied on NMDA current and was mediated by L-type calcium channels (Lo et al., 2002; Dilger et al., 2011; Dilger et al., 2015). In our experiments one large-amplitude EPSP could produce weak LTS without spikes or with one spike, a spike burst and short plateau potential. Suprathreshold activities, the spike bursts and plateau potentials, could also be mediated with L-type calcium channels and possible electrical contacts between thalamic cells (Lo et al., 2002; Lee et al., 2010; Dilger et al., 2011). Juxtacellular recordings confirmed the partial inactivation of spikes during plateau potential. Intrinsic AHP current (Fig.8) in VPM cells (Pirchio et al., 1997) could contribute to the intervals between spike bursts in thalamic cells and define the maximal frequency/occurrence of evoked bursts (low-pass filter). We also observed gamma-modulated spiking activity and the whole cell recordings revealed contribution of hyperpolarizing events in the generation of gamma oscillations. Gamma-modulated IPSPs could either decrease spike frequency/occurrence within bursts or prevent inactivation of fast sodium channels during plateau potential; as a result, VPM cells reproduced the rhythm of a presynaptic inhibitory cell. On average, gamma hot spots marked in spectrograms corresponded to the first, second and the third EPSP within each sensory response, that is, were locked to sensory inputs. Frequency/location of hot spots decreased with its ordinal (graded gamma activity). We observed graded gamma activity with silicone probes, juxtacellular recordings and patch clamp recordings in the thalamus. Sensory-evoked EPSPs in different VPM cells tended to form similar inter-EPSP intervals at the beginning of the response. Apparently, synchrony of independently-generated EPSPs in neighboring VPM cells decreased with EPSP ordinal which reduced firing rate in a corresponding inhibitory TRN cell/s and produced gradual decrease in gamma frequency. On the other hand, absence of the first hot spot, but presence of the second and/or the third indicates high synchrony of first EPSPs in neighboring VPM cells and following strong inhibitory feedback from the TRN at the beginning of a response. We propose that generation of graded gamma activity required intermediate synchrony of VPM cells. We also observed sensory-evoked graded gamma activity in the barrel cortex at the LFP level (needle LFP electrodes with small tips). We propose that sensory signals transmitted to the cortex by means of gamma activity. This is important to stress here that graded gamma activity and early gamma oscillations (Minlebaev et al., 2011; Yang et al., 2013) at the beginning of the response and detectable at the EEG level (Fig.9) corresponded to distinct levels of cortical synchrony. We recorded EEG signals with chlorinated silver wire (diameter 0.2 mm) and ~ 1 mm of the wire touched the cortical surface; thus, EEG gamma oscillations required synchrony within relatively large cortical area.

### Intrinsic neuronal oscillations and spindle-bursts in the immature thalamus

Immature thalamocortical cells in slices generate weak LTS and did not show intrinsic oscillatory properties (see Introduction). Intracellular pipettes and pipette solutions can change the intrinsic properties of small neurons with high input resistance (Tyzio et al., 2003) and noninvasive juxtacellular recordings are required for verification of weak intrinsic properties. Among our juxtacellular recordings, only one exhibited spontaneous bursting activity (Fig.6), and the bursts fulfilled the extracellular criteria of LTS bursts (Lenz et al., 1994; Jeanmonod et al., 1996). In contrast to immature thalamocortical cells, immature TRN cells in slices displayed relatively prominent intrinsic oscillatory properties (intrinsic LTS bursting) at room temperature 20-22°C (Wang et al., 2010), but intrathalamic TRN-dependent inhibition was weak in rodent pups compared to adult animals (Evrard and Ropert, 2009; Lee et al., 2010). In adult anesthetized mice, thalamic relay cells follow cortical active/UP states with one or several IPSPs without rebound LTS bursts and form 100-200 ms inter-event intervals (Sheroziya and Timofeev, 2014). In a shown example (Fig.9), spontaneous IPSPs and EPSPs in a VPM cell tended to form short-term oscillation-like patterns within gamma/beta frequency range. We detected neither spontaneous nor sensory-evoked sleep spindles in the VPM thalamic nucleus of rat pups.

Spontaneous and sensory-triggered alpha/beta oscillations (spindle-bursts) were firstly detected in the rodent somatosensory cortex (Khazipov et al., 2004) and then in visual (Hanganu et al., 2006; Colonnese and Khazipov, 2010; Kummer et al., 2016), motor (An et al., 2014) and prefrontal/prelimbic (Brockmann et al., 2011; Bitzenhofer et al., 2015) cortex in postnatal development. Spontaneous retinal waves trigger spindle-bursts in the neonatal visual cortex and removal of the retina reduces the frequency, but does not completely eliminate the cortical spindle-bursts (Hanganu et al., 2006). It has been also shown that subplate neurons and gap junctions are required for spindle-burst oscillations in newborn mice (Dupont et al., 2006). After selective removal of subplate neurons in the limb region of the primary somatosensory cortex, endogenous and sensory evoked spindle-bursts were largely abolished (Tolner et al., 2012). After developmental disappearance of the subplate by the end of the first postnatal week in mice the cortex generates NMDA-dependent spindle-bursts (Dupont et al., 2006). In the barrel cortex spontaneous and sensory-evoked spindle-bursts are composed of a negative delta wave nested with fluctuations within alpha/beta frequency range. Delta wave generation probably indicated depolarization of cortical cells and was sensitive to glutamatergic AMPA and NMDA receptor antagonists (Minlebaev et al., 2007, 2009). We observed similar sensory-evoked delta waves with nested oscillations in the barrel cortex. Sensory-evoked negative waves of smaller magnitude were also detected in the thalamus with silicon probes. Based on the intracellular recording of putative PO cell (Fig.9C), cortically generated spindle-bursts might represent either an early form of cortical active/UP states (Sanchez-Vives and McCormick, 2000; Steriade et al., 2001; Chauvette et al., 2010) or an early form of *slow* spindles (Timofeev and Chauvette, 2013). Besides other mechanisms, slow AHP currents in immature cortical neurons might underlie immature oscillation patterns (McCormick and Prince, 1987).

## Author contribution

MS performed the experiments and analyzed data, MS and RK designed the research project, MS and RK wrote the paper.

## Acknowledgements

We thank David Jappy for his helpful comments. We also thank Dr. Andrei Rozov for preparation of pipette solutions for patch-clamp recordings and Dr. Andrei Zakharov for technical assistance.

## References

An S, Kilb W, Luhmann HJ (2014) Sensory-evoked and spontaneous gamma and spindle bursts in neonatal rat motor cortex. J Neurosci 34:10870–10883.

Arcelli P, Frassoni C, Regondi MC, De Biasi S, Spreafico R (1997) GABAergic neurons in mammalian thalamus: a marker of thalamic complexity? Brain Res Bull 42:27–37.

Arsenault D, Zhang ZW (2006) Developmental remodelling of the lemniscal synapse in the ventral basal thalamus of the mouse. J Physiol-London 573:121–132.

Belford GR, Killackey HP (1979) The development of vibrissae representation in subcortical trigeminal centers of the neonatal rat. J Comp Neurol 188:63–74.

Bitzenhofer SH, Sieben K, Siebert KD, Spehr M, Hanganu-Opatz IL (2015) Oscillatory activity in developing prefrontal networks results from theta-gamma-modulated synaptic inputs. Cell Rep 11:486–497.

Blankenship AG, Feller MB (2010) Mechanisms underlying spontaneous patterned activity in developing neural circuits. Nat Rev Neurosci 11:18–29.

Brecht M, Sakmann B (2002) Whisker maps of neuronal subclasses of the rat ventral posterior medial thalamus, identified by whole-cell voltage recording and morphological reconstruction. J Physiol 538:495–515.

Brockmann MD, Poschel B, Cichon N, Hanganu-Opatz IL (2011) Coupled oscillations mediate directed interactions between prefrontal cortex and hippocampus of the neonatal rat. Neuron 71:332–347.

Buzsaki G, Mizuseki K (2014) The log-dynamic brain: how skewed distributions affect network operations. Nat Rev Neurosci 15:264–278.

Castro-Alamancos MA (2002) Properties of primary sensory (lemniscal) synapses in the ventrobasal thalamus and the relay of high-frequency sensory inputs. J Neurophysiol 87:946–953.

Chauvette S, Volgushev M, Timofeev I (2010) Origin of active states in local neocortical networks during slow sleep oscillation. Cereb Cortex 20:2660–2674.

Chen C, Regehr WG (2000) Developmental remodeling of the retinogeniculate synapse. Neuron 28:955–966.

Colonnese MT, Khazipov R (2010) “Slow activity transients” in infant rat visual cortex: a spreading synchronous oscillation patterned by retinal waves. J Neurosci 30:4325–4337.

Crunelli V, Cope DW, Hughes SW (2006) Thalamic T-type Ca2+ channels and NREM sleep. Cell Calcium 40:175–190.

Deschenes M, Timofeeva E, Lavallee P (2003) The relay of high-frequency sensory signals in the Whisker-to-barreloid pathway. J Neurosci 23:6778–6787.

Dilger EK, Shin HS, Guido W (2011) Requirements for synaptically evoked plateau potentials in relay cells of the dorsal lateral geniculate nucleus of the mouse. J Physiol 589:919–937.

Dilger EK, Krahe TE, Morhardt DR, Seabrook TA, Shin HS, Guido W (2015) Absence of plateau potentials in dLGN cells leads to a breakdown in retinogeniculate refinement. J Neurosci 35:3652–3662.

Dominguez S, Ma L, Yu H, Pouchelon G, Mayer C, Spyropoulos GD, Cea C, Buzsaki G, Fishell G, Khodagholy D, Gelinas JN (2021) A transient postnatal quiescent period precedes emergence of mature cortical dynamics. Elife 10.

Dupont E, Hanganu IL, Kilb W, Hirsch S, Luhmann HJ (2006) Rapid developmental switch in the mechanisms driving early cortical columnar networks. Nature 439:79–83.

Erzurumlu RS, Gaspar P (2012) Development and critical period plasticity of the barrel cortex. Eur J Neurosci 35:1540–1553.

Evrard A, Ropert N (2009) Early development of the thalamic inhibitory feedback loop in the primary somatosensory system of the newborn mice. J Neurosci 29:9930–9940.

Fanselow EE, Sameshima K, Baccala LA, Nicolelis MA (2001) Thalamic bursting in rats during different awake behavioral states. Proc Natl Acad Sci U S A 98:15330–15335.

Fernandez LMJ, Luthi A (2020) Sleep Spindles: Mechanisms and Functions. Physiol Rev 100:805–868.

Fogerson PM, Huguenard JR (2016) Tapping the Brakes: Cellular and Synaptic Mechanisms that Regulate Thalamic Oscillations. Neuron 92:687–704.

Ganmor E, Katz Y, Lampl I (2010) Intensity-dependent adaptation of cortical and thalamic neurons is controlled by brainstem circuits of the sensory pathway. Neuron 66:273–286.

Golshani P, Goncalves JT, Khoshkhoo S, Mostany R, Smirnakis S, Portera-Cailliau C (2009) Internally Mediated Developmental Desynchronization of Neocortical Network Activity. Journal of Neuroscience 29:10890–10899.

Groh A, Bokor H, Mease RA, Plattner VM, Hangya B, Stroh A, Deschenes M, Acsady L (2014) Convergence of cortical and sensory driver inputs on single thalamocortical cells. Cereb Cortex 24:3167–3179.

Guido W, Weyand T (1995) Burst responses in thalamic relay cells of the awake behaving cat. J Neurophysiol 74:1782–1786.

Hanganu IL, Ben-Ari Y, Khazipov R (2006) Retinal waves trigger spindle bursts in the neonatal rat visual cortex. J Neurosci 26:6728–6736.

Hirsch JC, Fourment A, Marc ME (1983) Sleep-related variations of membrane potential in the lateral geniculate body relay neurons of the cat. Brain Res 259:308–312.

Hwang K, Bertolero MA, Liu WB, D’Esposito M (2017) The Human Thalamus Is an Integrative Hub for Functional Brain Networks. J Neurosci 37:5594–5607.

Jager P, Moore G, Calpin P, Durmishi X, Salgarella I, Menage L, Kita Y, Wang Y, Kim DW, Blackshaw S, Schultz SR, Brickley S, Shimogori T, Delogu A (2021) Dual midbrain and forebrain origins of thalamic inhibitory interneurons. Elife 10.

Jahnsen H, Llinas R (1984) Ionic basis for the electro-responsiveness and oscillatory properties of guinea-pig thalamic neurones in vitro. J Physiol 349:227–247.

Jeanmonod D, Magnin M, Morel A (1996) Low-threshold calcium spike bursts in the human thalamus. Common physiopathology for sensory, motor and limbic positive symptoms. Brain 119 (Pt 2):363–375.

Khazipov R, Milh M (2018) Early patterns of activity in the developing cortex: Focus on the sensorimotor system. Semin Cell Dev Biol 76:120–129.

Khazipov R, Sirota A, Leinekugel X, Holmes GL, Ben-Ari Y, Buzsaki G (2004) Early motor activity drives spindle bursts in the developing somatosensory cortex. Nature 432:758–761.

Khazipov R, Zaynutdinova D, Ogievetsky E, Valeeva G, Mitrukhina O, Manent JB, Represa A (2015) Atlas of the Postnatal Rat Brain in Stereotaxic Coordinates. Front Neuroanat 9:161.

Kummer M, Kirmse K, Zhang C, Haueisen J, Witte OW, Holthoff K (2016) Column-like Ca(2+) clusters in the mouse neonatal neocortex revealed by three-dimensional two-photon Ca(2+) imaging in vivo. Neuroimage 138:64–75.

Lee SC, Cruikshank SJ, Connors BW (2010) Electrical and chemical synapses between relay neurons in developing thalamus. J Physiol 588:2403–2415.

Lenz FA, Kwan HC, Martin R, Tasker R, Richardson RT, Dostrovsky JO (1994) Characteristics of somatotopic organization and spontaneous neuronal activity in the region of the thalamic principal sensory nucleus in patients with spinal cord transection. J Neurophysiol 72:1570–1587.

Llinas RR, Steriade M (2006) Bursting of thalamic neurons and states of vigilance. J Neurophysiol 95:3297–3308.

Lo FS, Erzurumlu RS (2001) Neonatal deafferentation does not alter membrane properties of trigeminal nucleus principalis neurons. J Neurophysiol 85:1088–1096.

Lo FS, Guido W, Erzurumlu RS (1999) Electrophysiological properties and synaptic responses of cells in the trigeminal principal sensory nucleus of postnatal rats. J Neurophysiol 82:2765–2775.

Lo FS, Ziburkus J, Guido W (2002) Synaptic mechanisms regulating the activation of a Ca(2+)-mediated plateau potential in developing relay cells of the LGN. J Neurophysiol 87:1175–1185.

Luhmann HJ, Khazipov R (2018) Neuronal activity patterns in the developing barrel cortex. Neuroscience 368:256–267.

Marcano-Reik AJ, Blumberg MS (2008) The corpus callosum modulates spindle-burst activity within homotopic regions of somatosensory cortex in newborn rats. Eur J Neurosci 28:1457–1466.

Marlinski V, Beloozerova IN (2014) Burst firing of neurons in the thalamic reticular nucleus during locomotion. J Neurophysiol 112:181–192.

McCormick DA, Prince DA (1987) Post-natal development of electrophysiological properties of rat cerebral cortical pyramidal neurones. J Physiol 393:743–762.

McCormick DA, Trent F, Ramoa AS (1995) Postnatal development of synchronized network oscillations in the ferret dorsal lateral geniculate and perigeniculate nuclei. J Neurosci 15:5739–5752.

Mease RA, Sumser A, Sakmann B, Groh A (2016) Cortical Dependence of Whisker Responses in Posterior Medial Thalamus In Vivo. Cereb Cortex 26:3534–3543.

Minlebaev M, Ben-Ari Y, Khazipov R (2007) Network mechanisms of spindle-burst oscillations in the neonatal rat barrel cortex in vivo. J Neurophysiol 97:692–700.

Minlebaev M, Ben-Ari Y, Khazipov R (2009) NMDA receptors pattern early activity in the developing barrel cortex in vivo. Cereb Cortex 19:688–696.

Minlebaev M, Colonnese M, Tsintsadze T, Sirota A, Khazipov R (2011) Early gamma oscillations synchronize developing thalamus and cortex. Science 334:226–229.

Munoz A, Liu XB, Jones EG (1999) Development of metabotropic glutamate receptors from trigeminal nuclei to barrel cortex in postnatal mouse. J Comp Neurol 409:549–566.

Murata Y, Colonnese MT (2018) Thalamus Controls Development and Expression of Arousal States in Visual Cortex. J Neurosci 38:8772–8786.

Murata Y, Colonnese MT (2019) Thalamic inhibitory circuits and network activity development. Brain Res 1706:13–23.

Nunez A, Barrenechea C, Avendano C (1994) Spontaneous activity and responses to sensory stimulation in ventrobasal thalamic neurons in the rat: an in vivo intracellular recording and staining study. Somatosens Mot Res 11:89–98.

Panetsos F, Sanchez-Jimenez A (2010) Single unit oscillations in rat trigeminal nuclei and their control by the sensorimotor cortex. Neuroscience 169:893–905.

Perez Velazquez JL, Carlen PL (1996) Development of firing patterns and electrical properties in neurons of the rat ventrobasal thalamus. Brain Res Dev Brain Res 91:164–170.

Pinault D, Deschenes M (1992) The origin of rhythmic fast subthreshold depolarizations in thalamic relay cells of rats under urethane anaesthesia. Brain Res 595:295–300.

Pirchio M, Turner JP, Williams SR, Asprodini E, Crunelli V (1997) Postnatal development of membrane properties and delta oscillations in thalamocortical neurons of the cat dorsal lateral geniculate nucleus. J Neurosci 17:5428–5444.

Ramcharan EJ, Gnadt JW, Sherman SM (2000) Burst and tonic firing in thalamic cells of unanesthetized, behaving monkeys. Vis Neurosci 17:55–62.

Ramoa AS, McCormick DA (1994a) Developmental changes in electrophysiological properties of LGNd neurons during reorganization of retinogeniculate connections. The Journal of neuroscience : the official journal of the Society for Neuroscience 14:2089–2097.

Ramoa AS, McCormick DA (1994b) Enhanced Activation of Nmda Receptor Responses at the Immature Retinogeniculate Synapse. Journal of Neuroscience 14:2098–2105.

Sanchez-Vives MV, McCormick DA (2000) Cellular and network mechanisms of rhythmic recurrent activity in neocortex. Nat Neurosci 3:1027–1034.

Sandler VM, Puil E, Schwarz DW (1998) Intrinsic response properties of bursting neurons in the nucleus principalis trigemini of the gerbil. Neuroscience 83:891–904.

Sherman SM (2001) Tonic and burst firing: dual modes of thalamocortical relay. Trends Neurosci 24:122–126.

Sherman SM, Guillery RW (1996) Functional organization of thalamocortical relays. J Neurophysiol 76:1367–1395.

Sherman SM, Guillery RW (1998) On the actions that one nerve cell can have on another: distinguishing “drivers” from “modulators”. Proc Natl Acad Sci U S A 95:7121–7126.

Sheroziya M, Timofeev I (2014) Global intracellular slow-wave dynamics of the thalamocortical system. J Neurosci 34:8875–8893.

Sheroziya M, Timofeev I (2015) Moderate Cortical Cooling Eliminates Thalamocortical Silent States during Slow Oscillation. J Neurosci 35:13006–13019.

Steriade M, McCormick DA, Sejnowski TJ (1993) Thalamocortical oscillations in the sleeping and aroused brain. Science 262:679–685.

Steriade M, Timofeev I, Grenier F (2001) Natural waking and sleep states: a view from inside neocortical neurons. J Neurophysiol 85:1969–1985.

Takeuchi Y, Asano H, Katayama Y, Muragaki Y, Imoto K, Miyata M (2014) Large-scale somatotopic refinement via functional synapse elimination in the sensory thalamus of developing mice. J Neurosci 34:1258–1270.

Tennigkeit F, Schwarz DW, Puil E (1998) Postnatal development of signal generation in auditory thalamic neurons. Brain Res Dev Brain Res 109:255–263.

Timofeev I, Chauvette S (2013) The spindles: are they still thalamic? Sleep 36:825–826.

Tolner EA, Sheikh A, Yukin AY, Kaila K, Kanold PO (2012) Subplate neurons promote spindle bursts and thalamocortical patterning in the neonatal rat somatosensory cortex. J Neurosci 32:692–702.

Tyzio R, Ivanov A, Bernard C, Holmes GL, Ben-Ari Y, Khazipov R (2003) Membrane potential of CA3 hippocampal pyramidal cells during postnatal development. J Neurophysiol 90:2964–2972.

Urbain N, Fourcaud-Trocme N, Laheux S, Salin PA, Gentet LJ (2019) Brain-State-Dependent Modulation of Neuronal Firing and Membrane Potential Dynamics in the Somatosensory Thalamus during Natural Sleep. Cell Rep 26:1443–1457 e1445.

Urbain N, Salin PA, Libourel PA, Comte JC, Gentet LJ, Petersen CCH (2015) Whisking-Related Changes in Neuronal Firing and Membrane Potential Dynamics in the Somatosensory Thalamus of Awake Mice. Cell Rep 13:647–656.

Vitali I, Jabaudon D (2014) Synaptic biology of barrel cortex circuit assembly. Semin Cell Dev Biol 35:156–164.

Wang X, Yu G, Hou X, Zhou J, Yang B, Zhang L (2010) Rebound bursts in GABAergic neurons of the thalamic reticular nucleus in postnatal mice. Physiol Res 59:273–280.

Warren RA, Jones EG (1997) Maturation of neuronal form and function in a mouse thalamo-cortical circuit. J Neurosci 17:277–295.

Wei H, Bonjean M, Petry HM, Sejnowski TJ, Bickford ME (2011) Thalamic burst firing propensity: a comparison of the dorsal lateral geniculate and pulvinar nuclei in the tree shrew. J Neurosci 31:17287–17299.

Yamakado M (1985) [Postnatal development of barreloid neuropils in the ventrobasal complex of mouse thalamus: a histochemical study for cytochrome oxidase]. No To Shinkei 37:497–506.

Yang J-W, Hanganu-Opatz IL, Sun J-J, Luhmann HJ (2009) Three patterns of oscillatory activity differentially synchronize developing neocortical networks in vivo. The Journal of neuroscience : the official journal of the Society for Neuroscience 29:9011–9025.

Yang JW, An S, Sun JJ, Reyes-Puerta V, Kindler J, Berger T, Kilb W, Luhmann HJ (2013) Thalamic network oscillations synchronize ontogenetic columns in the newborn rat barrel cortex. Cereb Cortex 23:1299–1316.

